# The mechanical stability of Tension Gauge Tethers

**DOI:** 10.1101/2022.03.11.483943

**Authors:** Jingzhun Liu, Shimin Le, Mingxi Yao, Wenmao Huang, Zhikai Tio, Yu Zhou, Jie Yan

## Abstract

Mechanotransduction of cells relies on responding to tension transmitted along various supramolecular linkages. Tension gauge tethers (TGTs), short double-stranded DNA (dsDNA) fragments that undergo irreversible tension-dependent dissociation under shear-stretching mode, have been widely applied in live cell experiments to provide critical insights into the mechanotransduction activities of cells. However, the current physical understanding of the mechanical responses of TGTs remains limited, which restricts the range of information that can be extracted from experimental observations. In order to provide quantitative in-depth understanding and interpretation of experimental observations, in this work, we quantified the tension-dependent lifetime of TGTs from which the mechanical stability of TGTs under various physiologically relevant stretching conditions can be derived. Applications of the determined mechanical stability of TGTs to cell studies strongly suggest revisiting the previous interpretations of several reported experimental observations.

## I. INTRODUCTION

In cells, the actomyosin cytoskeleton network is linked to different cell membrane receptors and nuclear membrane proteins through sets of adaptor proteins. Some of the adhesive membrane receptors are further linked to extracellular matrix or neighboring cells, by which the cells in tissues are physically linked via cell-matrix and cell-cell adhesions [1–5]. Due to actomyosin contraction or external mechanical perturbations, dynamic tension is generated and transmitted along various intra/intercellular supramolecular linkages. By responding to the level of tension transmitted along such supramolecular linkages, cells are able to sense and respond to the mechanical cues in the local environment through processes referred to as mechanosensing and mechanotransduction, which play crucial roles in regulation of various cell behaviours including cell spreading, cell migration, cell differentiation, growth and tissue repair [1–10].

The mechanosensing and mechanotransduction functions of cells crucially depend on the mechanical properties of the tension transmission linkages. Different mechanical constraints applied to the cells result in different dynamic tensions on the linkages, driving tension-dependent conformational/structural changes of mechanosensing protein domains which in turn leads to tension-dependent interactions with binding factors. This way, the mechanical cues in the local environment can be converted into a downstream cascade of biochemical interactions [1, 11, 12]. Importantly, mounting evidences have suggested mechanical unfolding of mechanosensing protein domains as a major mechanism to expose cryptic binding sites for downstream signaling proteins [12–18]. A structural domain is associated with a critical tension *F*_c_ at which the domain has equal equilibrium unfolding and folding probabilities. *F*_c_ is typically in the order of a few piconewtons (pN) [16–20], which can be considered as the minimal tension needed to unfold the domains.

The tension-dependent conformational/structural transitions of the protein domains and the downstream interactions are time taking processes. Therefore, it is important to gauge the level of the tension and its duration on a tension-transmission supramolecular linkage. Correlating such information with cell behaviors, one can deduce the minimal requirements of the tension and duration needed for certain mechanosensing and mechanotranduction functions of cells. The recently developed tension sensors, short peptides or small biomolecular structures undergoing tension-dependent reversible conformational/structural changes, have made it possible to probe the tension ranges of several important tension-transmission linkages [21–28]. The physiologically relevant tension range has been reported in the order of a few pN [21, 23, 25, 28] on tension-transmission linkages at cell-matrix adhesions and cell-cell adherence junctions, coincident with the range of tensions needed to unfolding the mechanosensing protein domains in these linkages [14, 15, 17].

Tension Gauge Tethers (TGTs) [29], short double-stranded DNA (dsDNA) segments, has been inserted into tension-transmission linkages extracellularly at cell-matrix adhesions (Fig. 1a) and cell-cell adherence junctions, to probe the mechanical responses of the cells. In experiments, a TGT is inserted into a tension-transmission linkage via two attaching points, *P*_1_ at the end of one strand, and *P*_2_ on a position of the complementary strand separated from *P*_1_ by *M* base pairs (Fig. 1b). Its mechanical stability can be sensitively tuned by changing the distance of *P*_2_ from *P*_1_. When the DNA strands in a TGT are dissociated, the corresponding tension-transmission linkage is disconnected. Unlike tension sensors that undergo tension-dependent reversible conformational/structural changes, TGTs undergo irreversible tension-dependent strand dissociation, which is hereafter termed the rupturing transition of TGTs.

**Fig. 1.**
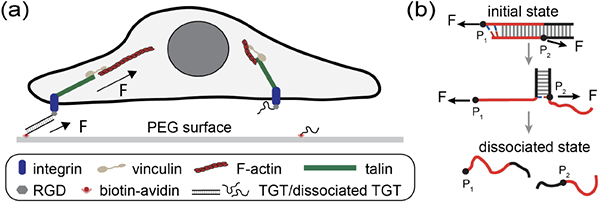
Illustration TGTs and applications in cell studies. (a) Schematics of a cell adhered to a surface mediated by integrins binding to RGDs tethered to the surface via TGTs. When the two strands in a TGT dissociate, the corresponding tension-transmission pathway disappears. (b) Illustration of the tension-dependent strand dissociation pathway of a TGT via strand-peeling from the ends [30]. The force attaching points, *P*_1_ and *P*_2_, are indicated by the black circles on the DNA strands.

Significant changes of cell behaviors in response to the changes of the stability of TGTs inserted at cell-matrix adhesions have been observed [29], indicating a direct impact of the TGT’s mechanical stability on the underlying mechanosensing and mechanotransduction activities of the tension-transmission linkages from integrin to the cytoskeleton network. Similarly, strong impacts of the TGT’s mechanical stability on the mechanosensing and mechanotransduction activities involved in cell-cell adherence junctions [26] and T-cell activation [31, 32] have been reported.

Despite the highly interesting experimental observations of the effects of TGT’s mechanical stability on cell’s mechanosensing and mechanotransduction activities, what insights that can be extracted from these observations remains unclear. TGTs were originally developed as extracellular tension sensors to report tension level based on a certain tension threshold termed the tension tolerance that is defined as the average tension needed for TGT’s rupturing within two seconds [29]. However, the tension-dependent irreversible rupturing of TGTs implies that a TGT can rupture at any tension, with a lifetime dependent on the tension. Therefore, without prior knowledge of the time scale of a tension duration, it is not straightforward to use TGTs to gauge the level of tension transmitted on a linkage [33]. To decode the information from the complex dependence of the cell’s mechanosensing and mechanotransduction activities on the mechanical stabilities of TGTs, a comprehensive understanding of the mechanical stabilities of TGTs is needed.

The current understanding of the mechanical stability of TGTs is vague. The applications of the quantity tension tolerance have faced several limitations. Firstly, the tension tolerance values of TGTs have not been experimentally calibrated. Instead, they have been estimated by deGennes’ formula derived based on static analysis of the elastic energy of the DNA [34], which does not consider the effects of temperature and the exact sequence of the TGTs. Secondly, due to the constrained time scale of two seconds, the concept of tension tolerance is a crude description of the mechanical stability of TGTs. It only provides a qualitative comparison of the mechanical stabilities of different TGTs -the higher the value of the tension tolerance, the greater the mechanical stability of a TGT. A complete description of the mechanical stability of TGTs is urgently needed to extract further information from experimental observations.

In this work, we experimentally calibrated several TGTs, including two that were frequently applied in previous cell studies, for tension-dependent lifetimes *τ* (*f*) at 37°C that is relevant to most cell studies. This quantity provides a comprehensive description of the mechanical stability of TGTs, from which the rupturing tensions or lifetimes of TGTs can be derived at various stretching conditions. Based on the results, the experimental observations from several previous cell studies were revisited for their interpretations.

## II. RESULTS

### A. Experimental design

We designed a single-molecule detector (Fig. 2a), consisting of a DNA hairpin construct spanned between two single-stranded DNA (ssDNA) handles, and an ssDNA oligo that is complementary to regions of both ssDNA handles adjacent to the fork of the hairpin marked by green and red, respectively. At tension below the threshold destablizing-tension of the hairpin (*F*_hairpin_ = 7.0 ± 0.7 pN at 37 °C), the hairpin remains stable, so that the green and red regions of the ssDNA oligo can hybridize with both ssDNA handles. The technical details of the magnetic tweezers used for the single-molecule study can be found in our previous publications [20, 35]. Further details on the manipulation of the single-molecule detector can be found in Supplemental Material I-IV [36].

**Fig. 2.**
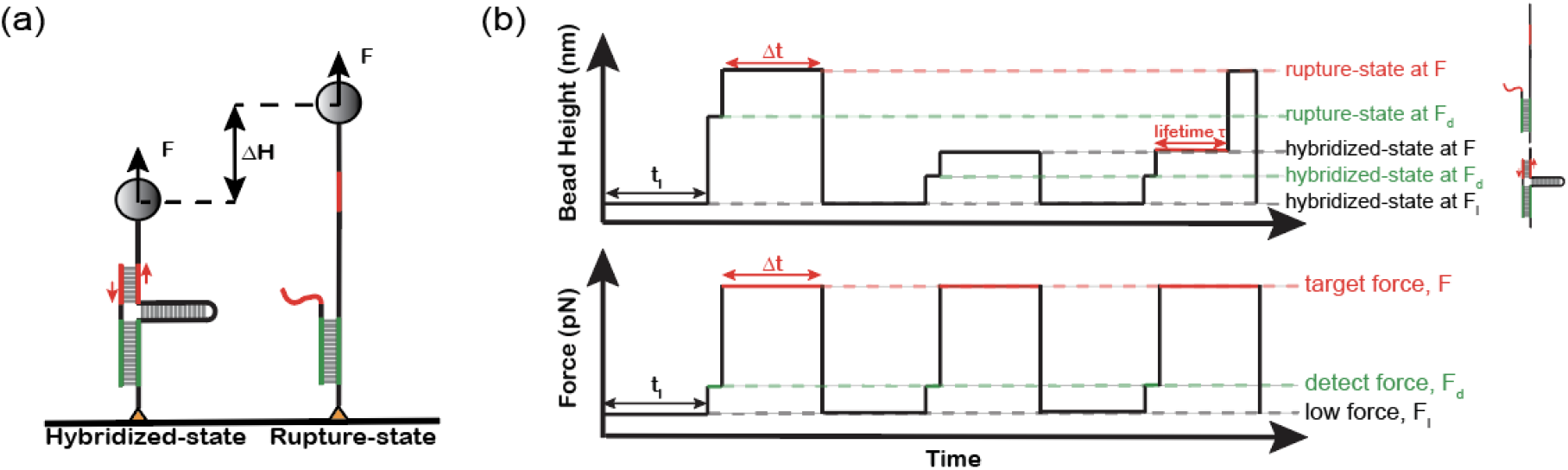
Tension-jump cycles to quantify the mechanical stability of a TGT. (a) Schematic of the designed DNA detector tethered between a glass surface and a superparamagnetic bead. The TGT is shown in red. The green DNA duplex is to anchor the ssDNA on the detector for higher experimental throughput. (b) Schematics of tension-jump cycle applied to measure the lifetime of the TGT. The state of the TGT is determined based on the bead height.

The red duplex corresponding to the region between *P*_1_ and *P*_2_ in a TGT is the target to be measured for its mechanical stability. At a tension *F*, the opening of the last base pair in the red duplex will place the hairpin under tension in an unzipping force geometry. This mimics an actual TGT, where the opening of the base-pair at the *P*_2_ position will render the duplex region after *P*_2_ in the same unzipping force geometry. At a tension greater than *F*_hairpin_, the opening of the last basepair in the red duplex is followed by immediate unzipping of the hairpin. Hence, the transition state of the designed construct is the same as that of actually applied TGTs (Fig. 1b, Supplemental Material V, VI)

When the red duplex is ruptured, it will lead to a large stepwise increase in the height of the bead that can be detected by our in-house constructed magnetic tweezers [20, 35]. The green duplex is made much longer than the red duplex to ensure that the rupturing transition only occurs on the red duplex (Supplemental Material VII [36]). After rupturing the red duplex, the ssDNA oligo remains stably bound to the detector via the green duplex (Fig. 2a, right), which allows the red duplex to quickly reform when tension is reduced to below *F*_hairpin_. This way, many data points can be obtained from one detector.

### B. The determination of *τ* (*f*) for TGTs

We quantified the tension-dependent rupturing probability of the red duplex over a time interval Δ*t* when it was held at a constant tension *F, p*_*F*_ (Δ*t*). The average lifetime of the red duplex *τ* (*f*) was obtained by fitting *p*_*F*_ (Δ*t*) data with a single-exponential function 1 *− e*^(−Δ*t/τ*)^. The *p*_*F*_ (Δ*t*) data were obtained by applying tension-jump cycles described in Figure 2b. Each tension-jump cycle starts with a tension, *F*_*l*_, at which the hairpin is stable, for one minute to allow the red duplex to form. In the subsequent step, tension is jumped to *F*_*d*_ which is slightly higher than the hairpin unzipping threshold tension, for two seconds to check whether the red duplex is formed. The two states of the red strand, hybridized or ruptured, can be unambiguously determined by their large extension difference (Fig. 2a). Then, the tension is jumped to a target higher tension F, for a holding time of Δ*t*, during which the red duplex may or may not rupture. Repeating the tension-jump cycles for multiple times, *p*_*F*_ (Δ*t*) is calculated as the number of cycles where the red duplex is ruptured, *N*_*r*_, divided by the total number of cycles, *N* (i.e., *p*_*F*_ (Δ*t*) = *N*_*r*_*/N*).

Most of cell studies have been performed at the human body temperature of 37 °C [29, 37–45]. As *τ* (*f*) of a TGT is highly sensitive to temperature (Supplemental Material VIII [36]), we quantified *τ* (*f*) of four TGTs (Table I) at 37 °C, including three (7-bp, 11-bp and 15-bp) widely used in previous studies and one modified (13-bp) from the 15-bp TGT. For each TGT, repeating the tension-jump cycles for multiple tethers (Supplemental Material IX [36]), the majority of the *p*_*F*_ (Δ*t*) data was determined from at least 50 cycles at each holding time. Figure 3a-c shows *p*_*F*_ (Δ*t*) obtained from three TGTs, a 15-bp TGT and an 11-bp TGT used in previous cell studies [29], as well as a 13-bp TGT generated by deleting the last two base pairs of the 15-bp TGT. The *p*_*F*_ (Δ*t*) curves of the three TGTs obtained at the same forces shift left as the lengths of TGTs decrease (Fig. 3d), suggesting decreased mechanical stability of shorter TGTs.

**TABLE I:**
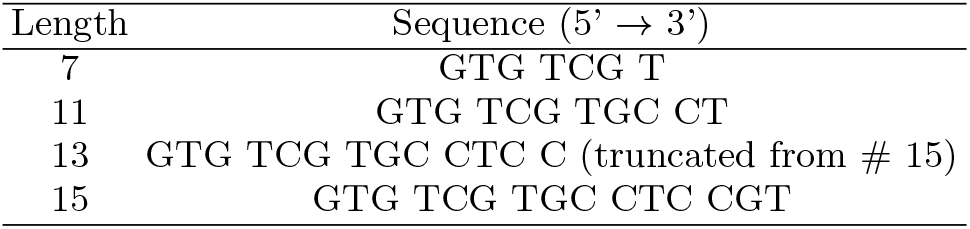
Sequences of TGTs [29]

**Fig. 3.**
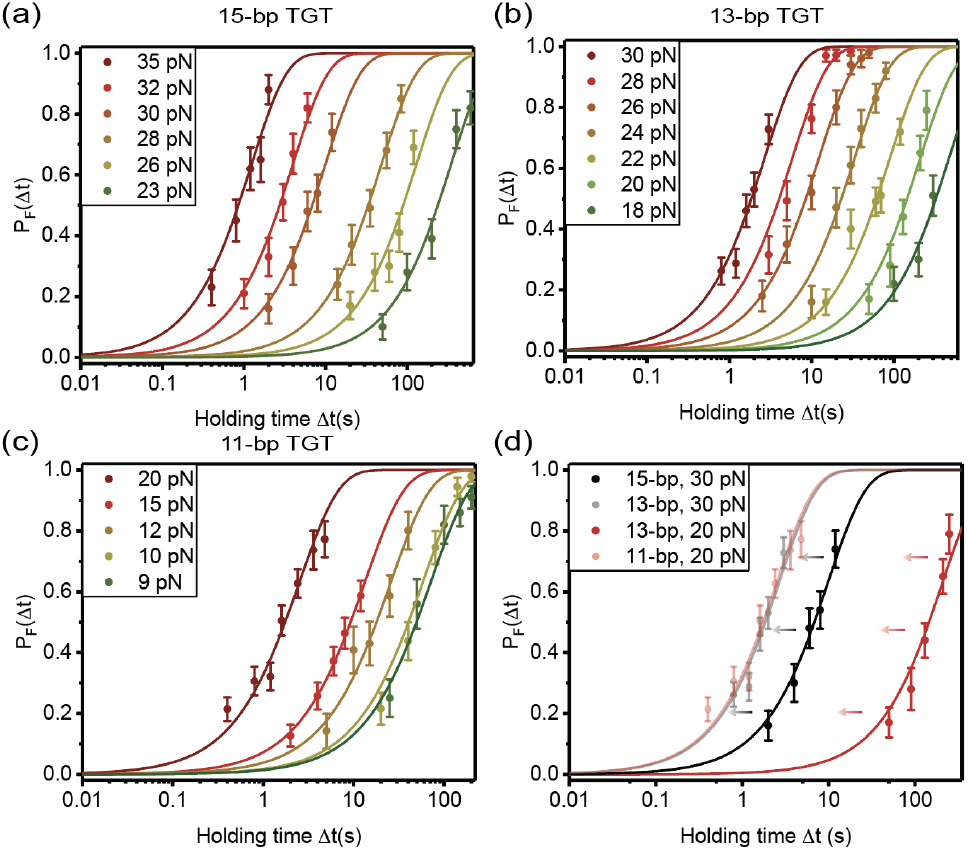
Tension-dependent lifetimes of three TGTs in Table I at 37 °C. The probabilities of 15-bp TGT (a), 13-bp truncated TGT (b), and 11-bp TGT (c) rupturing within different holding times were detected over a certain tension range. (d) Comparison of the *p*_*F*_ (Δ*t*) curves of 15-bp and 13-bp TGTs at 30 pN (dark and light black line) and 13-bp and 11-bp TGTs at 20 pN (dark and light red line). (mean ± standard error).

For each TGT, the best-fitted lifetime was obtained over a certain tension range (Fig. 4), with the errors obtained by bootstrap analysis (Supplemental Material X [36]). Outside the tension range, the lifetime of the TGT is either too short or too long to be accurately determined (Supplemental Material XI [36]). Therefore, *τ* (*f*) is extrapolated (solid line, Fig. 4) from the measured *τ* (*f*) data to tensions outside the range of measurement based on an expression derived from Arrhenius law of transition kinetics, 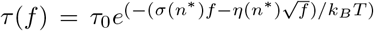, where *σ* = *n*^∗^(*l*_1*ss*_ *− l*_1*ds*_), and 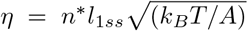 (Supplemental Material XII [36]). Here, *l*_1*ss*_ = 0.7 nm and *l*_1*ds*_ = 0.34 nm are the contour length of one nucleotide step of ssDNA and one base pair step of dsDNA, respectively, and *A* = 0.7 nm is the bending persistence length of ssDNA in 150 mM KCl [46, 47]. This model can be applied at tensions greater than a few pN where the worm-like chain polymer model of ssDNA is valid [48, 49].

**Fig. 4.**
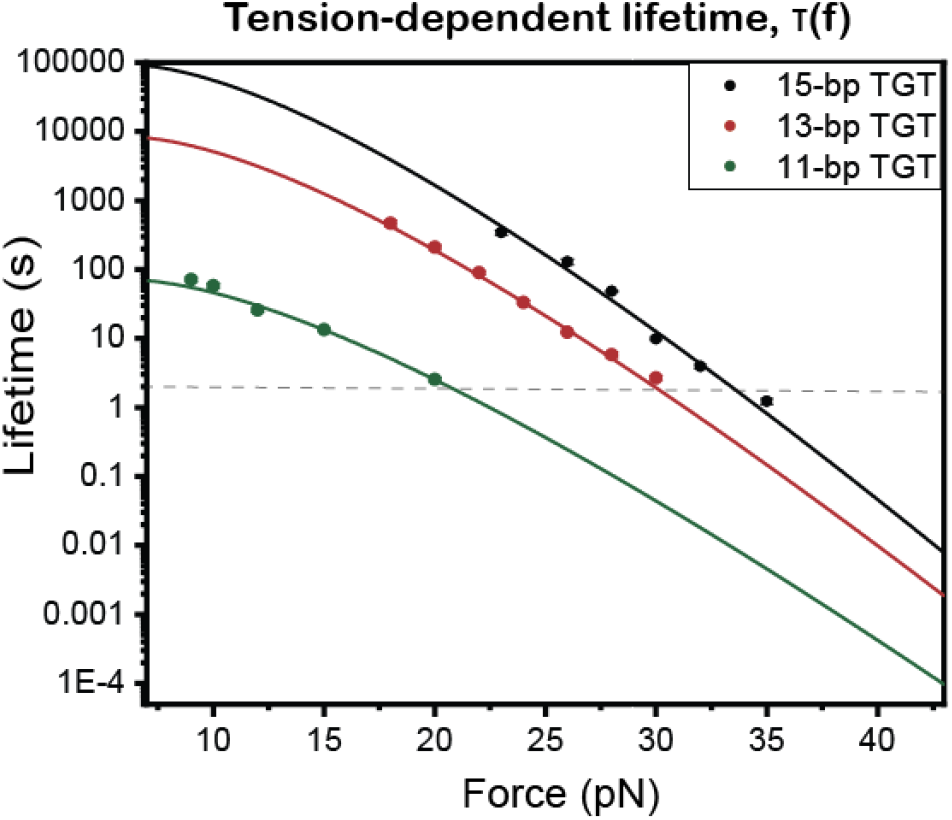
The mechanical stabilities of TGTs in Table I. The tension-dependent lifetime, *τ* (*F*), data is obtained from exponential fitting of measured *p*_*F*_ (Δ*t*) and is further extrapolated to tensions above a few pN using the structural-elastic model (Supplemental Material XII [36]).

### C. The tension tolerance and lifetime of TGT

We first applied the measured *τ* (*f*) to estimate the tension tolerances. The tension tolerance of a TGT has been defined as the tension needed to rupture the TGT over a time scale of 2 seconds. Our results (Fig. 4, dash line) show that at a lifetime of 2 seconds, the average rupturing tensions of the 11-bp, 13-bp and 15-bp TGTs are around 21 pN, 30 pN and 33 pN, respectively. Unexpectedly, these values are about 20 pN below the previously estimated values based on the deGennes’ formula derived from static analysis of the elastic energy of the DNA [34].

We then applied the measured *τ* (*f*) to estimate the life-times of TGTs at physiologically relevant tension range from a few pN to 15 pN of individual integrin-RGD complex [50–52]. The average lifetimes were determined to be 10^3^ *−* 10^4^ s for the 15-bp TGT, 10^2^ *−* 10^3^ s for the 13-bp TGT, and tens of seconds 10^1^ *−* 10^2^ s for the 11-bp TGT. *τ* (*f*) for the 7-bp TGT in Table I is too short to be determined (Supplemental Material XI [36]).

One can also derive the distribution of the lifetimes of a TGT based on *τ* (*f*) when the tension in a linkage changes with time. The distribution density function of the TGT’s lifetime can be expressed as 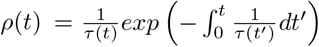, where *τ* ^−1^(*t*) is a time-varying rupturing rate of TGT caused by the tension *f* (*t*) that can be the time-varying. If the tension as a function of time is known, the lifetime distribution can be converted into a distribution of tensions at which the rupturing of TGT occurs. In a simple case when tension increases at a constant rate *r, f* (*t*) = *rt*, the TGT rupturing tension distribution is 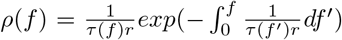 [53]. For the 11-bp TGT that provides the minimal stability to support the formation of nascent focal adhesions [29, 45], the peak TGT’s rupturing tension is in the range from 20 pN to 26 pN for loading rates in the order of a few pN/s (Fig. 5a, bottom panel) that are physiologically relevant (Supplemental Material XIII [36]).

**Fig. 5.**
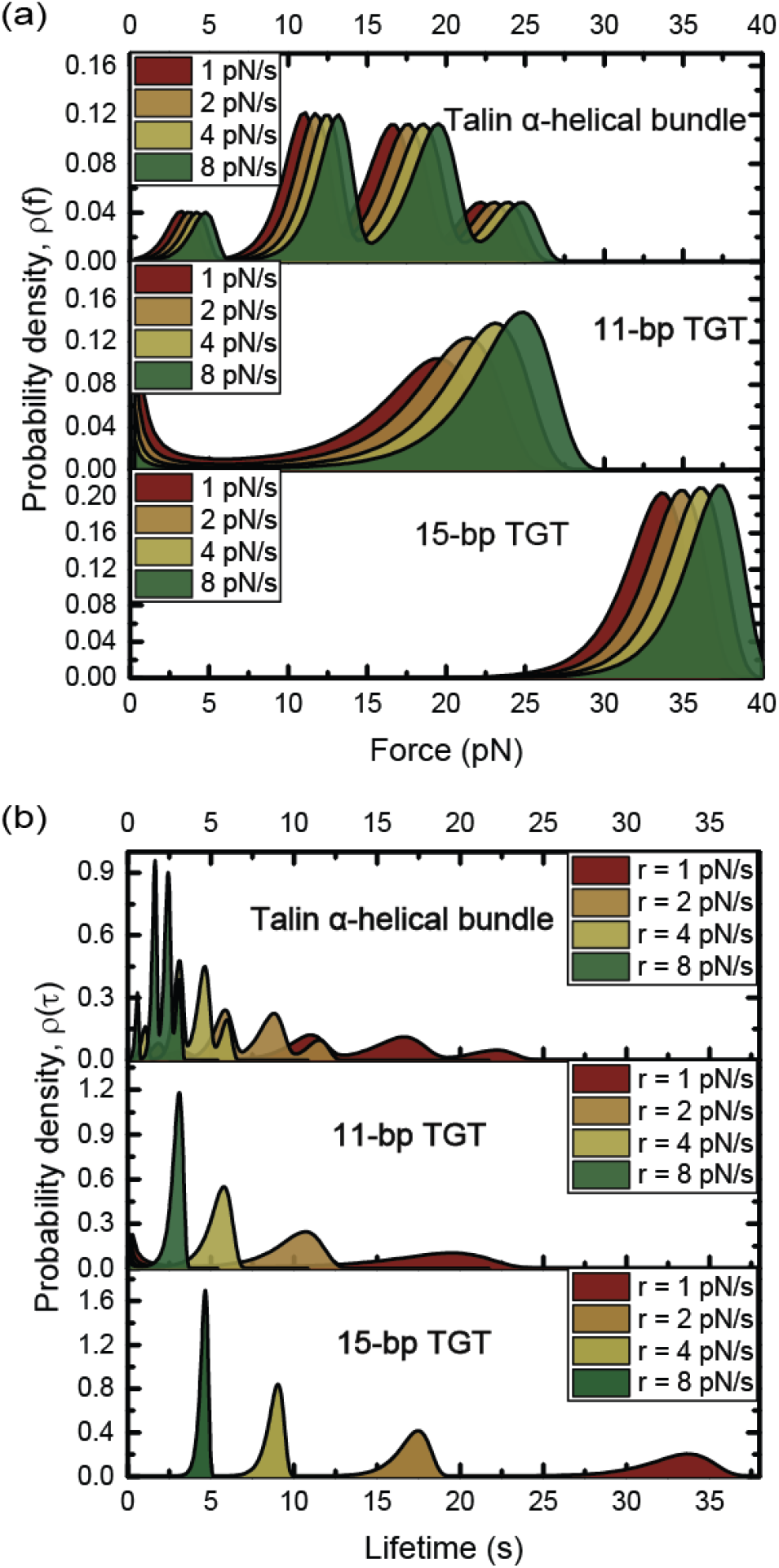
Comparison of the mechanical stabilities of talin *α*-helical bundles, 11-bp TGT, and 15-bp TGT. The rupturing tension probability densities (a) and the lifetime probability densities (b) of the *α*-helical bundles in talin rod (top panel), the 11-bp TGT (middle panel) and the 15-bp TGT (bottom panel) under increasing tension with different loading rates: 1, 2, 4, and 8 pN/s.

### D. Impacts of TGT’s mechanical stability on mechanotransduction

To support mechanotransduction, it requires that the TGTs can withstand high enough tension and provide long enough lifetime to activate the tension-bearing mechanosensing domains in the corresponding tension-transmission supramolecular linkage. In the case of talin/vinculin-mediated rigidity sensing that leads to the transition from nascent to matured focal adhesions, mechanical unfolding of the tension-bearing helical bundles in talin transducing tension between integrin and cytoskeleton and the subsequent binding and activation of vinculin are necessary [15, 16, 18, 54–57]. It can be reasoned that a newly assembled talin-mediated tension-transmission linkage in the nascent focal adhesions subjects the talin helical bundles to increasing tension toward the cell body due to the retrograde actin flow.

Figure 5b shows the comparison between the experimentally determined lifetime distribution of talin’s 13 tension-bearing helical bundles in the rod domain based on the tension-dependent unfolding rates measured in our previous study [18] and those for the 11-bp and 15-bp TGTs at the same loading rates *r* = 1, 2, 4, 8 pN/s. The 11-bp and 15-bp TGTs provide the minimal stability to support the formation of nascent and matured focal adhesions, respectively [29, 45]. The results show that the 11-bp TGT has a lifetime longer than majority of the talin *α*-helical bundles and the 15-bp TGT has a lifetime longer than all the talin *α*-helical bundles, at the corresponding loading rates. Over the loading rate range, the lifetimes (5—35 s) of the 15-bp TGT are significantly longer than those (3—20 s) of the 11-bp TGT. Since the tension is a monotonic function of time, the TGT’s and *α*-helical bundle’s lifetime distributions can also be converted into corresponding rupturing/unfolding tension distributions. The results show that the rupturing tensions of both 11-bp and 15-bp TGTs are larger than the unfolding tensions of talin *α*-helical bundles, and that the 15-bp TGT’s peak rupturing tension is more than 10 pN greater than that of the 11-bp TGT (Fig. 5a).

## III. DISCUSSIONS

In summary, we have measured the tension-dependent lifetimes *τ* (*f*) of several TGTs at 37°C over a lifetime range from a few seconds to several hundreds of seconds, which covers a major range of time scale involved in tension-dependent conformational/structural changes of mechanonsensing protein domains and tension-dependent interactions of signaling proteins with the protein domains in tension-transmission linkages. Based on a physical model of tension-dependent kinetics of unfolding or rupturing of biomolecular complexes [46], *τ* (*f*) is further extrapolated to time scale outside the range. As *τ* (*f*) is a complete description of the mechanical stability of TGTs, the knowledge of *τ* (*f*) will be highly useful to decode the information from cell studies using TGTs.

A direct impact of our study is that it leads to corrections to the tension tolerance values of TGTs estimated previously. Our results show that at 37°C, the average tensions to rupture the TGTs within two seconds are around 21 pN for the 11-bp TGT, 30 pN for the 13-bp TGT and 33 pN for the 15-bp TGT. The 11-bp and 15-bp TGTs have been widely used in cell studies [29, 37, 44, 45, 58–60]. Their tension tolerance values were previously estimated to be 43 pN and 50 pN, respectively, which are 22 pN and 17 pN higher than our measured values. Our results also lead to release of a constraint of time scale for using tension tolerance to report whether tension in a tension-transmission linkage exceeds a certain threshold. The previous definition of tension tolerance is the average tension at which a TGT has an average lifetime of two seconds, which restricts the applications of the TGTs to tension transmission durations in the order of seconds. Using *τ* (*f*), one can use TGTs to report whether tension exceeds certain threshold for tension-transmission durations that are not restricted to two seconds.

Using tension tolerance to estimate tension in a linkage requires prior knowledge of the tension duration. A TGT of a higher tension tolerance may rupture under conditions associated with a longer tension duration, whereas a TGT of a lower tension tolerance may persist under conditions associated with a shorter tension duration. Therefore, unless the tension durations are comparable, one cannot determine relative tensions between different stretching conditions. For an example, in the original paper by Wang et al. [29], it was reported that 11-bp TGT provided the minimal stability to support initial cell spread and adhesion, while more stable longer TGTs were needed to form matured focal adhesions. Consistently, a more recent study observed only small nascent focal adhesions appeared in cells seeded on 11-bp TGT, whereas much longer (*>*18 bp) TGTs were required to form larger matured focal adhesions [45]. For both studies, the observations were interpreted as that higher tension is applied to the intgerin-RGD complexes at matured focal adhesions than that at nascent focal adhesions. However, as an initially assembled tension-transmission linkage from integrin to cytoskeleton at nascent focal adhesions could have a significantly shorter linkage lifetime than that at matured focal adhesions, the tension durations across TGTs between the two conditions can be very different. It is well possible that the TGTs may be exposed to higher tension for shorter duration at the nascent focal adhesions, and lower tension for a longer duration at matured focal adhesions, which could also explain the observations. As alternative interpretations exist, it is insufficient to determine the relative tension levels at different cell spreading stages or different locations on the cells solely based on the signals from TGTs.

We reason that *τ* (*f*) can provide information beyond tension. The reciprocal of the tension-dependent lifetime of TGTs, *τ* ^−1^(*f*), has a meaning of the rupturing probability per unit of time. It reflects an important fact that thermal fluctuation can spontaneously rupture a TGT at any tension within any time interval as pointed previously [33]. Therefore, based on *τ* (*f*), TGTs are promising to inform us the time scale of tension transmission in a mechanosensing supramolecular linkage needed for specific processes. At nascent focal adhesions, the tension transmitted on integrin was measured to be from a few pN to around 13 pN using reversible tension sensors [24]. According to the measured *τ* (*f*), at tensions in 10-15 pN, the lifetimes of the 11-bp TGT are 10-100 seconds. At lower tensions, the TGT’s lifetime should be even longer. Interestingly, the lifetimes of the 11-bp TGT over this force range are comparable to the average lifetime of nascent focal adhesions, which were measured to tens of seconds [61]. This result provides an explanation why the 11-bp TGT is sufficient for the formation of nascent focal adhesions.

Based on *τ* (*f*), TGTs may provide further insights into the time scale of tension transmission in a mechanosensing supramolecular linkage needed for proper mechanotransduction. Using the integrin/talin/vinculin-mediated rigidity sensing for an example, mechanical unfolding of talin rod domains that in turn recruits and activates vinculin is necessary for the initial rigidity sensing of the substrate that leads to maturation of focal adhesions on sufficiently rigid substrate surface [1, 12]. A newly assembled talin-mediated mechanosensing linkage is likely subjected to increasing tension due to the retrograde actin flow [62]. Figure 5b shows that over physiologically relevant loading rates of a few pN per second, the lifetimes of the 11-bp TGT are in the order of seconds. According to Figure 5b, the lifetime of the 11-bp TGT is long enough to unfold most talin rod domains. However, since the 11-bp TGT cannot support the transition from nascent to matured focal adhesions [29, 45], it is likely that the time scale might not be sufficiently long to support the downstream mechanotransduction that requires recruiting and activating vinculin. A more stable TGT, such as the 15-bp TGT that was reported able to support the formation of matured focal adhesions [45], is needed to provide a longer lifetime (Fig. 5b) not only for unfolding the talin rod domains but also for the downstream mechanotransduction steps.

A limitation of the current study is that only the part of the TGTs in the shear-stretching mode is considered, which restricts the transition state structure corresponds to that the last base pair in the shear-mode part of duplex is ruptured. This imposes that the tensions should be higher than the base pair destabilizing threshold under the unzipping geometry (*>* 10 pN) for the remaining part of the DNA to apply this model. A complete understanding of the tension-dependent lifetime of TGTs including the lower tension range (*<* 10 pN) is important, which is an ongoing project in our group.

## ACKNOWLEDGEMENTS

We thank Alexander Bershadsky and Pakorn Tony Kanchanawong for critical reading of the manuscript. The research was funded by the Singapore Ministry of Education Academic Research Funds Tier 2 (MOE2019-T2-1-099, MOE-T2EP50220-0015) and the Ministry of Education under the Research Centres of Excellence programme.

## AUTHOR CONTRIBUTIONS

J.L. and J.Y. designed the research, interpreted the experimental data and wrote the manuscript. J.L. and Z.T., carried out the single-molecule experiments and performed data analysis. J.L., S.L., M.Y., Y.Z., and W.H. contributed to the design and synthesized the DNA detector for single-molecule experiments. J.Y. supervised the research.

## Notes

### Competing Interest Statement

The authors have declared no competing interest.

### Summary of Updates

The manuscript is revised to include 1) the comparison of the lifetimes of TGTs and the rod domains in talin proteins under physiological range of loading rates; 2) extended discussions on the information of tensions and tension durations that can be extracted from TGTs; 3) more in-depth discussions of the TGT's tension-dependent lifetime measurement on the interpretations of previous TGT-based cell studies.

## References

[1] T. Iskratsch, H. Wolfenson, and M. P. Sheetz, Appreciating force and shape—the rise of mechanotransduction in cell biology, Nature reviews Molecular cell biology 15, 825 (2014).

[2] C. Uhler and G. Shivashankar, Regulation of genome organization and gene expression by nuclear mechanotransduction, Nature reviews Molecular cell biology 18, 717 (2017).

[3] K. H. Vining and D. J. Mooney, Mechanical forces direct stem cell behaviour in development and regeneration, Nature reviews Molecular cell biology 18, 728 (2017).

[4] T. Panciera, L. Azzolin, M. Cordenonsi, and S. Piccolo, Mechanobiology of yap and taz in physiology and disease, Nature reviews Molecular cell biology 18, 758 (2017).

[5] B. Ladoux and R.-M. Mège, Mechanobiology of collective cell behaviours, Nature reviews Molecular cell biology 18, 743 (2017).

[6] I. Schoen, B. L. Pruitt, and V. Vogel, The yin-yang of rigidity sensing: how forces and mechanical properties regulate the cellular response to materials, Annual Review of Materials Research 43, 589 (2013).

[7] Z. Sun, S. S. Guo, and R. Fässler, Integrin-mediated mechanotransduction, Journal of Cell Biology 215, 445 (2016).

[8] F. Martino, A. R. Perestrelo, V. Vinarský, S. Pagliari, and G. Forte, Cellular mechanotransduction: from tension to function, Frontiers in physiology 9, 824 (2018).

[9] A. Angulo-Urarte, T. van der Wal, and S. Huveneers, Cell-cell junctions as sensors and transducers of mechanical forces, Biochimica et Biophysica Acta (BBA)-Biomembranes 1862, 183316 (2020).

[10] M. Michael and M. Parsons, New perspectives on integrin-dependent adhesions, Current opinion in cell biology 63, 31 (2020).

[11] Y. Wang, J. Yan, and B. T. Goult, Force-dependent binding constants, Biochemistry 58, 4696 (2019).

[12] B. T. Goult, N. H. Brown, and M. A. Schwartz, Talin in mechanotransduction and mechanomemory at a glance, Journal of Cell Science 134, jcs258749 (2021).

[13] F. Nakamura, T. M. Osborn, C. A. Hartemink, J. H. Hartwig, and T. P. Stossel, Structural basis of filamin a functions, The Journal of cell biology 179, 1011 (2007).

[14] S. M. Pang, S. Le, A. V. Kwiatkowski, and J. Yan, Mechanical stability of αt-catenin and its activation by force for vinculin binding, Molecular biology of the cell 30, 1930 (2019).

[15] M. Yao, B. T. Goult, H. Chen, P. Cong, M. P. Sheetz, and J. Yan, Mechanical activation of vinculin binding to talin locks talin in an unfolded conformation, Scientific reports 4, 1 (2014).

[16] A. Del Rio, R. Perez-Jimenez, R. Liu, P. Roca-Cusachs, J. M. Fernandez, and M. P. Sheetz, Stretching single talin rod molecules activates vinculin binding, Science 323, 638 (2009).

[17] M. Yao, W. Qiu, R. Liu, A. K. Efremov, P. Cong, R. Seddiki, M. Payre, C. T. Lim, B. Ladoux, R.-M. Mege, et al., Force-dependent conformational switch of α-catenin controls vinculin binding, Nature communications 5, 1 (2014).

[18] M. Yao, B. T. Goult, B. Klapholz, X. Hu, C. P. Toseland, Y. Guo, P. Cong, M. P. Sheetz, and J. Yan, The mechanical response of talin, Nature communications 7, 1 (2016).

[19] M. Schlierf, F. Berkemeier, and M. Rief, Direct observation of active protein folding using lock-in force spectroscopy, Biophysical journal 93, 3989 (2007).

[20] H. Chen, H. Fu, X. Zhu, P. Cong, F. Nakamura, and J. Yan, Improved high-force magnetic tweezers for stretching and refolding of proteins and short dna, Biophysical Journal 100, 517 (2011).

[21] C. Grashoff, B. D. Hoffman, M. D. Brenner, R. Zhou, M. Parsons, M. T. Yang, M. A. McLean, S. G. Sligar, C. S. Chen, T. Ha, et al., Measuring mechanical tension across vinculin reveals regulation of focal adhesion dynamics, Nature 466, 263 (2010).

[22] M. Morimatsu, A. H. Mekhdjian, A. S. Adhikari, and A. R. Dunn, Molecular tension sensors report forces generated by single integrin molecules in living cells, Nano letters 13, 3985 (2013).

[23] B. L. Blakely, C. E. Dumelin, B. Trappmann, L. M. McGregor, C. K. Choi, P. C. Anthony, V. K. Duesterberg, B. M. Baker, S. M. Block, D. R. Liu, et al., A dna-based molecular probe for optically reporting cellular traction forces, Nature methods 11, 1229 (2014).

[24] Y. Zhang, C. Ge, C. Zhu, and K. Salaita, Dna-based digital tension probes reveal integrin forces during early cell adhesion, Nature communications 5, 1 (2014).

[25] T.-J. Kim, S. Zheng, J. Sun, I. Muhamed, J. Wu, L. Lei, X. Kong, D. E. Leckband, and Y. Wang, Dynamic visualization of α-catenin reveals rapid, reversible conformation switching between tension states, Current biology 25, 218 (2015).

[26] X. Wang, Z. Rahil, I. T. Li, F. Chowdhury, D. E. Leckband, Y. R. Chemla, and T. Ha, Constructing modular and universal single molecule tension sensor using protein g to study mechano-sensitive receptors, Scientific reports 6, 1 (2016).

[27] P. Roca-Cusachs, V. Conte, and X. Trepat, Quantifying forces in cell biology, Nature cell biology 19, 742 (2017).

[28] P. Ringer, A. Weißl, A.-L. Cost, A. Freikamp, B. Sabass, A. Mehlich, M. Tramier, M. Rief, and C. Grashoff, Multiplexing molecular tension sensors reveals piconewton force gradient across talin-1, Nature methods 14, 1090 (2017).

[29] X. Wang and T. Ha, Defining single molecular forces required to activate integrin and notch signaling, Science 340, 991 (2013).

[30] S. Cocco, J. Yan, J.-F. Léger, D. Chatenay, and J. F. Marko, Overstretching and force-driven strand separation of double-helix dna, Physical Review E 70, 011910 (2004).

[31] Y. Liu, L. Blanchfield, V. P.-Y. Ma, R. Andargachew, K. Galior, Z. Liu, B. Evavold, and K. Salaita, Dna-based nanoparticle tension sensors reveal that t-cell receptors transmit defined pn forces to their antigens for enhanced fidelity, Proceedings of the National Academy of Sciences 113, 5610 (2016).

[32] V. P.-Y. Ma, Y. Hu, A. V. Kellner, J. M. Brockman, A. Velusamy, A. T. Blanchard, B. D. Evavold, R. Alon, and K. Salaita, The magnitude of lfa-1/icam-1 forces finetune tcr-triggered t cell activation, Science advances 8, eabg4485 (2022).

[33] M. Mosayebi, A. A. Louis, J. P. Doye, and T. E. Ouldridge, Force-induced rupture of a dna duplex: from fundamentals to force sensors, ACS nano 9, 11993 (2015).

[34] Pierre-Gilles, de, and Gennes, Maximum pull out force on dna hybrids, Comptes Rendus De Lacadémie Des Sciences (2001).

[35] X. Zhao, X. Zeng, C. Lu, and J. Yan, Studying the mechanical responses of proteins using magnetic tweezers, Nanotechnology 28, 414002 (2017).

[36] See Supplemental Material at [URL will be inserted by publisher] for detailed information of magnetic-tweezer experiment, base pair-destablizing tension, temperature-sensitive mechanical stability of TGTs, bootstrap sampling methods, structural-elastic model, and the physiologically relevant tension range.

[37] F. Chowdhury, I. T. Li, B. J. Leslie, S. Doğanay, R. Singh, X. Wang, J. Seong, S.-H. Lee, S. Park, N. Wang, et al., Single molecular force across single integrins dictates cell spreading, Integrative Biology 7, 1265 (2015).

[38] M. H. Jo, W. T. Cottle, and T. Ha, Real-time measurement of molecular tension during cell adhesion and migration using multiplexed differential analysis of tension gauge tethers, ACS Biomaterials Science & Engineering 5, 3856 (2018).

[39] Y. Zhang, Y. Qiu, A. T. Blanchard, Y. Chang, J. M. Brockman, V. P.-Y. Ma, W. A. Lam, and K. Salaita, Platelet integrins exhibit anisotropic mechanosensing and harness piconewton forces to mediate platelet aggregation, Proceedings of the National Academy of Sciences 115, 325 (2018).

[40] Y. Zhao, A. Sarkar, and X. Wang, Peptide nucleic acid based tension sensor for cellular force imaging with strong dnase resistance, Biosensors and Bioelectronics 150, 111959 (2020).

[41] Y. Zhao, Y. Wang, A. Sarkar, and X. Wang, Keratocytes generate high integrin tension at the trailing edge to mediate rear de-adhesion during rapid cell migration, Iscience 9, 502 (2018).

[42] Y. Zhao, K. Pal, Y. Tu, and X. Wang, Cellular force nanoscopy with 50 nm resolution based on integrin molecular tension imaging and localization, Journal of the American Chemical Society 142, 6930 (2020).

[43] Y. Hu, V. P.-Y. Ma, R. Ma, W. Chen, Y. Duan, R. Glazier, B. G. Petrich, R. Li, and K. Salaita, Dna-based microparticle tension sensors (μts) for measuring cell mechanics in non-planar geometries and for highthroughput quantification, Angewandte Chemie 133, 18192 (2021).

[44] H. Li, C. Zhang, Y. Hu, P. Liu, F. Sun, W. Chen, X. Zhang, J. Ma, W. Wang, L. Wang, et al., A reversible shearing dna probe for visualizing mechanically strong receptors in living cells, Nature Cell Biology 23, 642 (2021).

[45] C.-Y. C. Chien, S.-H. Chou, and H.-H. Lee, Integrin molecular tension required for focal adhesion maturation and yap nuclear translocation, Biochemistry and Biophysics Reports 31, 101287 (2022).

[46] S. Guo, Q. Tang, M. Yao, H. You, C. H. Le, S., and J. Yan, Structural-elastic determination of the force-dependent transition rate of biomolecules, Chemical science 9, 5871 (2018).

[47] X. Zhang, H. Chen, S. Le, I. Rouzina, P. S. Doyle, and J. Yan, Revealing the competition between peeled ssdna, melting bubbles, and s-dna during dna overstretching by single-molecule calorimetry, Proceedings of the National Academy of Sciences 110, 3865 (2013).

[48] H. Fu, S. Le, C. Hu, K. Muniyappa, and Y. Jie, Force and atp hydrolysis dependent regulation of reca nucleoprotein filament by single-stranded dna binding protein, Nucleic Acids Research, 924 (2013).

[49] A. Bosco, J. Camunas-Soler, and F. Ritort, Elastic properties and secondary structure formation of single-stranded dna at monovalent and divalent salt conditions, Nucleic Acids Research 42, 2064 (2013).

[50] F. Li, S. D. Redick, H. P. Erickson, and V. T. Moy, Force measurements of the α5β1 integrin–fibronectin interaction, Biophysical Journal 84, 1252 (2003).

[51] E. Kokkoli, S. E. Ochsenhirt, and M. Tirrell, Collective and single-molecule interactions of α5β1 integrins, Langmuir the Acs Journal of Surfaces & Colloids 20, 2397 (2004).

[52] E. L. Baker and M. H. Zaman, The biomechanical integrin, Journal of Biomechanics 43, 38 (2010).

[53] O. K. Dudko, G. Hummer, and A. Szabo, Intrinsic rates and activation free energies from single-molecule pulling experiments, Physical review letters 96, 108101 (2006).

[54] Y. Wang, M. Yao, K. B. Baker, R. E. Gough, and J. Yan, Force-dependent interactions between talin and full-length vinculin, (2021).

[55] P. Kanchanawong, G. Shtengel, A. M. Pasapera, E. B. Ramko, M. W. Davidson, H. F. Hess, and C. M. Waterman, Nanoscale architecture of integrin-based cell adhesions, Nature 468, 580 (2010).

[56] I. Thievessen, P. M. Thompson, S. Berlemont, K. M. Plevock, S. V. Plotnikov, A. Zemljic-Harpf, R. S. Ross, M. W. Davidson, G. Danuser, S. L. Campbell, et al., Vinculin–actin interaction couples actin retrograde flow to focal adhesions, but is dispensable for focal adhesion growth, Journal of Cell Biology 202, 163 (2013).

[57] J. L. Bays and K. A. DeMali, Vinculin in cell–cell and cell–matrix adhesions, Cellular and Molecular Life Sciences 74, 2999 (2017).

[58] Z. Wan, X. Chen, H. Chen, Q. Ji, Y. Chen, J. Wang, Y. Cao, F. Wang, J. Lou, Z. Tang, et al., The activation of igm-or isotype-switched igg-and ige-bcr exhibits distinct mechanical force sensitivity and threshold, Elife 4, e06925 (2015).

[59] X. Wang, J. Sun, Q. Xu, F. Chowdhury, M. Roein-Peikar, Y. Wang, and T. Ha, Integrin molecular tension within motile focal adhesions, Biophysical journal 109, 2259 (2015).

[60] F. Chowdhury, S. Doğanay, B. J. Leslie, R. Singh, K. Amar, B. Talluri, S. Park, N. Wang, and T. Ha, Cdc42-dependent modulation of rigidity sensing and cell spreading in tumor repopulating cells, Biochemical and biophysical research communications 500, 557 (2018).

[61] C. K. Choi, M. Vicente-Manzanares, J. Zareno, L. A. Whitmore, A. Mogilner, and A. R. Horwitz, Actin and α-actinin orchestrate the assembly and maturation of nascent adhesions in a myosin ii motor-independent manner, Nature cell biology 10, 1039 (2008).

[62] I. Andreu, B. Falcones, S. Hurst, N. Chahare, X. Quiroga, A.-L. Le Roux, Z. Kechagia, A. E. Beedle, A. Elosegui-Artola, X. Trepat, et al., The force loading rate drives cell mechanosensing through both reinforcement and cytoskeletal softening, Nature communications 12, 1 (2021).

