## Supplementary Information for "The mechanical stability of Tension Gauge Tethers"

### Supplemental Material

#### I. Single-molecule constructs

All DNA oligos used in this study were custom synthesized by Integrated DNA Technologies (Coralville, Iowa) shown in Table SI. The DNA-based detector with two dsDNA handles and a hairpin region was synthesized following the standard PCR, digestion, annealing, ligation, and purification protocols.

#### II. Single-molecule manipulation

The single-molecule manipulation experiments were conducted using an in-house-made back-scattered vertical magnetic tweezers with a spatial resolution of 1 nm and temporal resolution of 200 Hz. DNA-based detector was tethered to the coverslip through Biotin/Traptavidin linkage, while the other end of the detector was linked to a superparamagnetic bead through Thiol/Epoxy reaction. This system was performed in a laminar flow channel. The extension change of the tethered DNA was measured based on the height change of the superparamagnetic beads tethered to the DNA-based detector under force. The details of the force calibration and control for the single-molecule magnetic tweezers experiments have been described in previous papers [1, 2].

Experiments were performed in a standard solution containing: 1 × PBS and 1% BSA at multiple temperatures (23 (room temperature), 30, and 37 °C). To achieve temperatures higher than room temperature, a non-magnetic nanocarbon-based thermal film (Ultra-Thin Flexible Heater, Pelonis Technologies) was applied on the top of the sample channel.

TABLE SI: DNA sequences. Underscore indicates the DNA segments that form the TGT region under shearing force geometry.

| Oligo ID | Sequence (5' → 3') |
|---|---|
| Com_TGT_7 bp | ACG ACA CTT TTC GAG TCT GTG CAC AAG CAG CGT GGA GGT ATG ACA ACC ACG GAA TGC ATT TTT CTG GCA GCG GGC TTC ATA TTC TGT GTG CTT ATG CTT GCC GAC ATG GGA (100 bp anchor) |
| Com_TGT_11 bp | A GGC ACG ACA CTT TTC GAG TCT GTG CAC AAG CAG CGT GGA GGT ATG ACA ACC ACG GAA TGC ATT TTT CTG GCA GCG GGC TTC ATA TTC TGT GTG CTT ATG CTT GCC GAC ATG GGA |
| Com_TGT_13 bp | GGA GGC ACG ACA CTT TTC GAG TCT GTG CAC AAG CAG CGT GGA GGT ATG ACA ACC ACG GAA TGC ATT TTT CTG GCA GCG GGC TTC ATA TTC TGT GTG CTT ATG CTT GCC GAC ATG GGA |
| Com_TGT_15 bp | AC GGA GGC ACG ACA CTT TTC GAG TCT GTG CAC AAG CAG CGT GGA GGT ATG ACA ACC ACG GAA TGC ATT TTT CTG GCA GCG GGC TTC ATA TTC TGT GTG CTT ATG CTT GCC GAC ATG GGA |
| Hairpin_TGT_30 bp | /5Phos/CTT GTG CAC AGA CTC GTT TAAT ATA CTA ATA TCT TAT ATT AAC TAT TAT TTTT ATA ATA GTT AAT ATA AGA TAT TAG TAT ATT TTT GTG TCG TGC CTC CGT CGAC |
| Handle1_1st_BstXI_CAGC | TTAACCAGCAGCGTGGAGGTATGACAACCACGGAATG |
| Handle1_2nd_2bio | /52-Biosg/ TCAGTACGCTACGGCAAATG |
| Handle2_1st_BstXI_CGAC | TTAACCAGCGACGTGG TCAGTACGCTACGGCAAATG |
| Hande2_2nd_BstXI_ATCC | TTAACCAGATCCGTGG AGGTATGACAACCACGGAATG |
| 3lable_BstXI_ATCC | TCTAGCTCTTCAAGCATCC |
| 3lable_BstXI_3Thiol | /5Phos/GCT TGA AGA GCT AGA / 3ThioMC3-D/ |
| Flank1_TGT | /5Phos/ACG GAG GCA CGA CAC |
| Flank2 | CGAGTCTGTGCACAAGCAGC |

##### III. Force calibration of superparamagnetic microbead

For a given pair of magnets, at the same magnet-bead distance $d$, the ratio of the forces applied to two different superparamagnetic beads, $\frac{F_2(d)}{F_1(d)} = c_{21}$, is a constant which does not depend on $d$ [1, 2]. Therefore, if a standard force-distance curve, $F_1(d)$, is calibrated for a bead '1' over a wide range of force, then the force on another bead '2' can be directly extrapolated by $F_2(d) = c_{21}F_1(d)$, where the value of $c_{21}$ can be calibrated at any convenient magnet-bead distance $d$.

The standard curve $F_1(d)$ was obtained by measuring drift speed $\nu$ of a superparamagnetic bead in glycerol solution with known viscosity $\eta_{gly} = 1.412$ Pa·s at given bead-magnet distances by the equation $F_1(d) = 6\pi\eta_{gly}R\nu$, where $R = 1.4 \pm 0.5$ $\mu$m (mean ± s.t.d) (CV < 3 %) is the bead radius. In our experiment, the value of $c_{21}$ was calibrated at a particular magnet-bead distance at midst of the overstretching of dsDNA occurs in 1× PSB, corresponding to a force of 65±2 pN [3]. Therefore, the uncertainly in the force-calibration due to the bead size variation and the determination of $c_{21}$ contributed to 5 % relative error (mainly due to the bead size variation).

##### IV. Bead height change upon the TGT rupture

The hairpins we used in this study to generate all the data in the main text have a 30-bp hairpin stem with 4-nt terminal loop (see Supplemental Material II: DNA sequences, Hairpin_TGT_30 bp). A longer hairpin (200-bp stem and a 4-nt terminal loop) was used to generate the data to demonstrate the temperature sensitivity in Supplemental Material VIII. This longer hairpin construct was used in the early stage of our study. Later we switched to the 30-bp hairpin design because it is easier to make the construct. What is important is that different hairpin lengths will not cause a change to the results, as they all share the same transition state corresponding to opening the last base pair in the shear-force region of TGTs.

When the TGT is ruptured, the concurrent unzipping of the DNA hairpin at a certain force $F$ lead to a large stepwise bead height increase, which is the same as the extension change under the force, $\Delta x(F)$. For the 30-bp+4-nt hairpin, the collective data of the extension change obtained at 40 pN from 10 independent detectors are analysed, $\Delta x(40 \text{ pN}) = 38.5 \pm 2.1$ nm. When the DNA hairpin is fully unzipped, in total 64 nt of ssDNA is released under tension. The extension of the ssDNA can be calculated based on the force-extension curve predicted with worm-like chain polymer model, $\chi_{ss}(F) = L_{ss}(1 - \sqrt{\frac{k_BT}{4A_{ss}F}})$, where $L_{ss} = 64 \times 0.7$ nm= 44.8 nm is the contour length of a 64 nt ssDNA and $A_{ss} = 0.7$ nm is the persistence length of ssDNA [4, 5]. The estimated extension of the ssDNA, $\chi_{ss}(40 \text{ pN}) = 35.8$ nm, is reasonably consistent with the measured extension change of 38.5 nm, suggesting that the observed stepwise bead height increase is the fully unzipping of the DNA hairpin. Similarly, for the 200-bp+4-nt hairpin, 404-nt of ssDNA will be released under tension. The extension of the ssDNA is also calculated $\chi_{ss}(40 \text{ pN}) = 228.6$ nm, which is reasonably consistent with the measured extension change of $\Delta x(40 \text{ pN}) = 237.2 \pm 8.3$ nm obtained at 40 pN from 10 independent detectors are analysed.

TABLE SII: DNA base pair energy and base pair-destabilizing tension under zipping and shearing force geometry. The base pair energy and base pair-destabilizing tension of ten independent dinucleotide sequences are calculated at typical room temperature of 24 °C and physiologically relative temperature of 37 °C.

| | $\varepsilon_i(k_BT)$ | | $F_{bp,shear}(pN)$ | |
|---|---|---|---|---|
| | 24 °C | 37 °C | 24 °C | 37 °C |
| AA | 1.7506 | 1.2686 | 50 | 44 |
| AT | 1.645 | 1.2008 | 48 | 42 |
| TA | 1.2136 | 0.7505 | 41 | 34 |
| CA | 2.6407 | 2.1482 | 65 | 59 |
| GT | 2.6231 | 2.1369 | 64 | 59 |
| CT | 2.3258 | 1.869 | 59 | 55 |
| GA | 2.3962 | 1.9142 | 60 | 55 |
| CG | 3.873 | 3.2861 | 84 | 78 |
| GG | 3.1759 | 2.7422 | 73 | 70 |
| GC | 3.924 | 3.3959 | 85 | 80 |

#### V. Design of TGTs, the stretching geometry, and their strand-dissociation transition pathways

The TGTs are originally designed as short dsDNA segments with two force attaching points [6], $P_1$ on the end of one strand and $P_2$ on the opposite strand M base pairs from $P_1$, on a dsDNA segment of total number of base pairs $N = 21$ (Fig. S1(a)). $P_1$ and $P_2$ will divide a TGT into two regions: $R_s$ refers to DNA segment between $P_1$ and $P_2$ with base pairs in this region under shearing geometry and $R_u$ refers to the rest part of TGT that under unzipping geometry. The predominant strand-dissociation transition is a sequential breakage of the base pairs from the $P_1$ end, which defines a transition pathway indicated by the position of the fork (i.e., the number of broken base pairs) (Fig. S1(b)). When the fork position is between $P_1$ and $P_2$, a shearing force geometry is applied. When the fork position passes $P_2$, the force geometry switches to unzipping. This leads to a transition state precisely located at $P_2$. Our design of the single-molecule construct has a similar transition pathways and the same transition state as the actual TGTs (Fig. S1(c)).

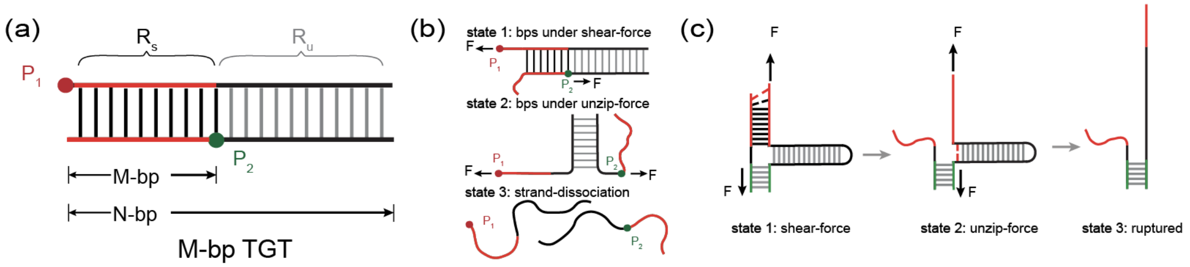


FIG. S1. Illustration of the design of TGTs (a), the strand-dissociation transition pathways of a 11-bp TGT (b), and that of our single-molecule detectors (c).

#### VI. The energy landscapes of our detectors and the actual TGTs.

The strand-dissociation process of a TGT can be understood based on a random walk of the fork position, with the forward (base pair opening) and backward (base pair formation) rates dependent on the tension. Opening a base pair at the fork results in elimination of a dsDNA base pair step under tension and the production of two ssDNA nucleotide steps, one of which is placed under the same tension. The energy cost of the opening of $i^{th}$ base pair from the ends is $\Delta g_i(F) = \varepsilon_i + (\phi_{ss}(F) - \phi_{ds}(F))$, where $\varepsilon_i$ is the zero-tension base pair opening energy cost. $\phi_{ss}(F) = \int_0^F -x_{ss}(f)df$ and $\phi_{ds}(F) = \int_0^F -x_{ds}(f)df$ are the tension-dependent conformation free energies of 1-nt of ssDNA step and 1-bp dsDNA base pair step, respectively, where $x_{ss}(f)$ and $x_{ds}(f)$ are the corresponding force-extension curves of 1-nt ssDNA and 1-bp dsDNA. The $i^{th}$ base pair-destabilizing tension is determined by the equation $\Delta g_i(F_{bp,i}) = 0$. The value of $\varepsilon_i$ depends on ten independent dinucleotide sequences of the nearest-neighboring base pairs [7]. At 37 °C, the base pair energy $\varepsilon_i$ is in a range of (0.75 $k_BT$, 3.4 $k_BT$), corresponding to a range of $F_{bp,i} \in$(35 pN, 80 pN) ($F_{bp,shear}$ in Table SII).

Similarly, for DNA segment under unzipping force geometry, opening a base pair at the fork results in elimination of a dsDNA base pair step and the production of two ssDNA nucleotide steps that are placed under the same tension. In this case, the energy cost of the opening of $i^{th}$ base pair from the ends is $\Delta g_i(F) = \varepsilon_i + 2\phi_{ss}(F))$. Both our designed single-molecule detecter and the original designed TGTs are DNA segments that first undergo shear-stretch and then undergo unzip-stretch geometry. In this case, the energy landscape of our detecter and TGTs under different stretching force can be calculated (Fig. S2), where $n$ is noted as the state where DNA strand dissociate from one end to the $n$th base pair. It is clear that our designed single-molecule detector has very similar energy landscape as the actual TGT over the force range from 0 to 70 pN. And our detector shares the same energy barrier with the actual TGT when the force is above the unzipping critical force of the $R_u$ region. Based on Arrhenius law of chemical reaction rate, the tension-dependent lifetime quantified using our detector should be close to that of the actual TGT with forces larger than the hairpin unzipping critical force. In Figure S2, we also provide energy landscapes of the strand-dissociation transition of TGTs based on sequence-dependent nearest-neighboring basepair energy with the basepair energy data from the work by Felix Ritort [8]. They also predict very similar energy barrier for all the tested TGTs.

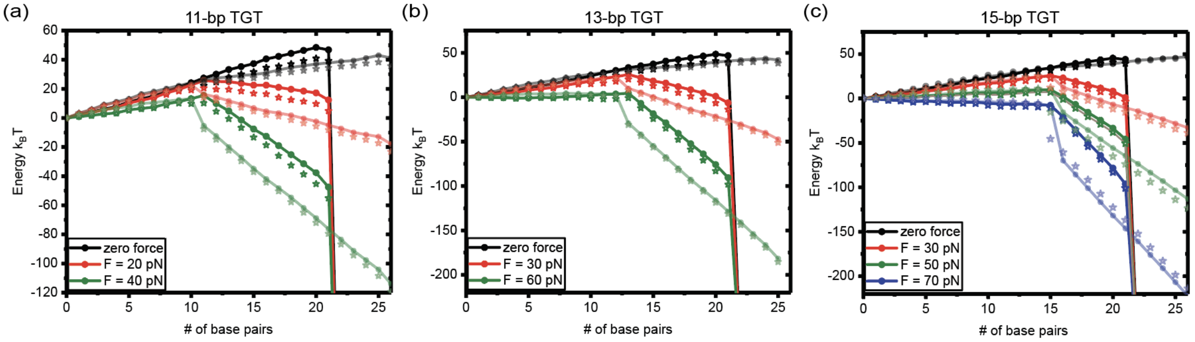


FIG. S2. Comparison of energy landscapes of our designed construct (deep color) and that of the actual TGTs (light color) with 11, 13, and 15-bp under shearing geometry and more than 6-bp under unzipping geometry with the unified nearest-neighboring base pair energy data [7] (dot) and that with the base pair energy data from the work by Felix Ritort [8] (star). The stretching force increases from top to bottom. Our designed construct produces highly similar energy landscape to those of the actual TGTs and shares the same energy barrier and transition states (n = 15) at forces higher than the unzipping critical force for the unzip region.

##### VII. The negligible possibility of the rupturing of the green duplex in experiments

At forces below min $F_{bp,i}$, a condition satisfied in most experiments, the transition state of the strand dissociation corresponds to a structure where the two strands are held by the last base pair. For a TGT with N base pairs, the corresponding energy barrier is $\Delta G^*(F) = \sum_{(i=1)}^{N} \Delta g_i(F)$. Therefore, besides the dependence on the sequence, $\Delta G^*(F)$ increases as the length increases. The lifetime of a TGT under a force can be obtained by Arrhenius law, $\tau(F) = \tau_0 e^{\Delta G^*(F)/k_B T}$, where the pre-factor $\tau_0$ is positively dependent on the number of base pair $N$ due to a longer diffusion distance till reaching the transition state according to the Kramers kinetic theory [9]. Together, $\tau(F)$ can be sensitively tuned by varying the sequence and the length of TGTs. The red duplex (i.e., the TGT sequence) and green duplex (to anchor the TGT on the single-molecule detector) have similar GC percentage. The green duplex was designed to have 80 more base pairs. As discussed above, this ensures that the strand dissociation transition observed in the experiments occurs only on the red duplex due to the drastically increased energy barrier.

##### VIII. Temperature-sensitive mechanical stability of TGTs

Considering that the stability of each DNA base pair depends on temperature [10], the rate of strand dissociation of a TGT, which requires rupturing multiple base pairs until reaching the transition state, may be sensitive to temperature changes. To probe the sensitivity of the tension-dependent lifetime of TGTs to temperature, we quantified the mechanical stability of a 16 base pairs (bp) TGT at $40 \pm 2$ pN at 23 °C and 30 °C, using the approach described in the main text. The probability of the strand dissociation as a function of the holding time was determined at the two temperatures (Fig. S3). From the data the best-fit average lifetimes of the 16-bp TGT at 23 °C and 30 °C, were determined to be $148.09 \pm 8.88$ s and $1.02 \pm 0.07$ s, respectively, where the errors were obtained by bootstrap analysis (Supplemental Material VII). The average lifetime for this 16-bp TGT at 40 pN and a higher temperature of 37 °C was too short ($\ll$0.3 s) to be determined by our instrument. Together, these results indicate highly sensitive temperature dependence of TGTs. Hence, to apply TGTs in cell studies, the TGTs must be calibrated at experimentally relevant temperatures.

##### IX. Data from individual tethers

As shown in previous section, $\tau(f)$ of a TGT is highly sensitive to temperature, we quantified $\tau(f)$ of TGTs (Table I) at 37 °C, including three (7-bp, 11-bp and 15-bp) widely used in previous studies and one modified (13-bp) from the 15-bp TGT. For each TGT, repeating the tension-jump cycles for multiple tethers ($N = 15$ for 15-bp TGT, $N = 13$ for 13-bp TGT, and $N = 11$ for 11-bp TGT), the $p_F(\Delta t)$ data were determined from at least 50 cycles at each holding time. Data for each force condition were obtained from 3-6 independent tethers. Figure S4 shows $p_F(\Delta t)$ obtained from individual tethers of three TGTs, a 15-bp TGT and an 11-bp TGT used in previous cell studies [6], as well as a

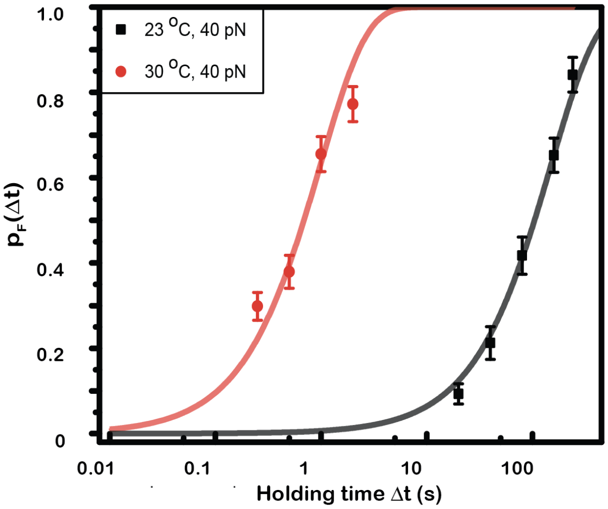


FIG. S3. Temperature-sensitive mechanical stability of a 16-bp TGT (3' CTA CTG ACC TGG CTG C 5'). Rupturing probability of the TGT within certain holding times $\Delta t$ =0.25 s,0.5 s,1 s,2 s,20 s,40 s,80 s,160 s,240 s, and the single exponential decay fitting: $p_F(\Delta(t) = 1 - e^{(-\Delta t/\tau)}$, from 10 independent DNA detectors. The error bars were obtained by bootstrap analysis (Supplemental Material VIII).

13-bp TGT generated by deleting the last two base pairs of the 15-bp TGT.

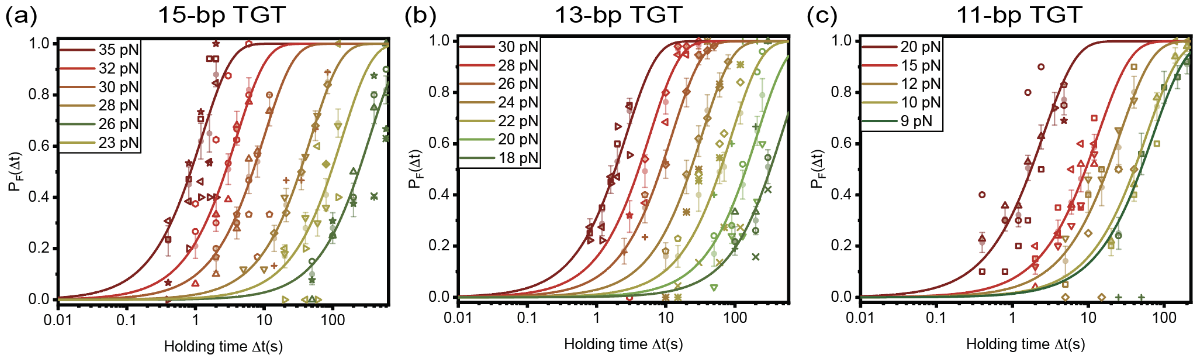


FIG. S4. $p_F(\Delta t)$ obtained from individual tethers of three TGTs, a 15-bp TGT and an 11-bp TGT and their single exponential decay fitting: $p_F(\Delta t) = 1 - e^{(-\Delta t/\tau)}$. Different symbols indicate data points obtained from different tethers (circle, square, rhombus, triangle-up, triangle-down, triangle-left, triangle-right, pentagon, hexagon, star, cross, diagonal-cross, double-cross, dash, and cross-square). The error bars were obtained by bootstrap analysis (Supplemental Material VIII).

##### X. The bootstrap sampling methods

Bootstrap is a resampling method that generate groups of independent sample data from an existing sample data with the same data size. From our approach, we obtained $N_r$, which is the number of cycles that a TGT ruptured corresponding to a bead height change at constant target force, $F$, during a certain time interval, $\Delta t$, after repeating force jump cycles for $N$ times. Thus, $p_F(\Delta t) = \frac{N_r}{N}$.

The resampling procedure are as followed: Assuming the data pool containing $N_r$ rupturing events and $N - N_r$ non-rupturing events ($N$ events in total), we randomly choose one event from $N$ events, record whether it is a rupturing event or a non-rupturing event, and then put it back to the data pool. Following this step and choose $N$ times, we obtain a group of $N$ resampled data with $N_r^b$ to be rupturing events, resulting to a resampled $p_F^b(\Delta t) = \frac{N_r^b}{N}$. Repeating this resampling process for 100 times, we will have 100 groups of resampled data and thus have 100 independent

$p_F^b(\Delta t, i) = \frac{N_r^b(i)}{N}(i = 1, 2, \dots, 100)$. The mean value and the standard deviation of the rupturing probability under a certain condition $(F, T, \Delta t)$ can be calculated from $p_F^b(\Delta t, i)(i = 1, 2, \dots, 100)$. Then, with resampled $p_F^b(\Delta t, i)$ of a range of $\Delta t$, by the single exponential decay fitting, the average lifetime, $\tau_F$, and its standard error can be found.

#### XI. Limited experimental force range

For the 15-bp TGT, at forces greater than 35 pN, the lifetime became too short to be measured due to limited sampling rate (200 Hz) and the limited speed of force change (force jump took 0.3 s to complete). At forces below 23 pN, the lifetime exceeded 300 s, making it time consuming to obtain enough data to calculate $p_F(\Delta t)$. For the 13-bp and 11-bp TGT, the experimental force range in this study were 18-30 pN and 9-20 pN. We further examined the mechanical stability of a 7-bp TGT, which was not stable and only sustained for seconds at physiological force range at 23 °C. When the temperature increased to 37 °C, we found that its lifetime was ≪0.3 s at physiological force range and was not detectable using our instrument.

#### XII. Structural-elastic model of force-dependent unfolding rate

At low force, the differential force-dependent entropic extension fluctuation between the native and the transition states leads to the force dependence of the transition rate. This is related to the differential structural-elastic properties of molecules between their native state and the transition state [11]. At forces larger than $\frac{k_BT}{b^0}$, $\frac{k_BT}{b^*}$, and $\frac{k_BT}{A}$, this rate has a simple asymptotic expression: $k(F) = k_0 e^{\frac{\sigma F + \sigma F^2/2 - \eta F^{1/2}}{k_BT}}$, which contains a kinetic parameter $k_0$ and three model parameters $\sigma = L^* + b^* - b^0 - (\frac{k_BT}{\gamma^*} - \frac{k_BT}{\gamma^0})$, $\alpha = \frac{b^*}{\gamma^*} - \frac{b^0}{\gamma^0}$, and $\eta = L^* \sqrt{\frac{k_BT}{A}}$ referring to Fig. S5.

In the case of the force-dependent strand dissociation of TGTs, $\gamma^0 = \gamma^* \in (1000, 1500)$ pN [12], are the stretching rigidity of dsDNA, which makes the term $\alpha F^2/2$ negligible compared with the other two terms. As the two remaining model parameter $\sigma$ and $\eta$ only determined by the transition state, they can be described by a single parameter $n^*$, which is the number of ruptured base pairs at the transition state: $\sigma = n^*(l_{1ss} - l_{1ds})$ and $\eta = n^* l_{1ss} \sqrt{\frac{k_BT}{A}}$, where $l_{1ss}$ and $l_{1ds}$ are the contour length of per ssDNA nucleotide and dsDNA base pair.

In this study, the structural-elastice model is applied for the tension-dependent strand dissociation of TGTs. Denoting $n^*$ the number of ruptured base pairs at the transition state, it can be shown that $\tau(f) = \tau_0 e^{(-(\sigma(n^*)f - \eta(n^*)\sqrt{f})/k_BT)}$. Here, $l_{1ss} = 0.7$ nm and $l_{1ds} = 0.34$ nm are the contour length of one nucleotide step of ssDNA and one base pair step of dsDNA, respectively, and $A = 0.7$ nm is the bending persistence length of ssDNA in 150 mM KCl [11, 13]. By fitting the measured data with this model, the transition states ($n^*$) of these TGTs under shearing tension geometry can be obtained.

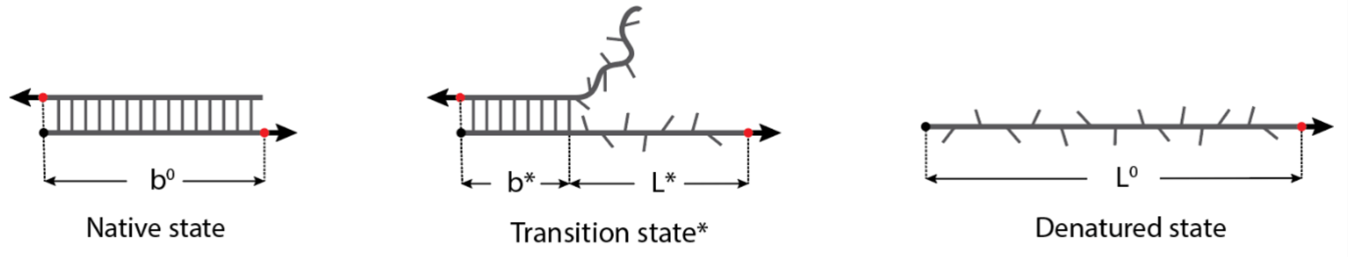


FIG. S5. Illustration of the native, the transition, and the denatured state of a dsDNA under shear force (transition occurs from one-open end of dsDNA). The native state is a hybridized dsDNA, with a length of $b^0$ and stretching rigidity of $\gamma^0$. The transition state consists of a segment of hybridized dsDNA with a length of $b^*$ and stretching rigidity of $\gamma^*$ as well as a segment of dissociated ssDNA with a contour length of $L^*$. The denatured state is a fully dissociated ssDNA with a contour length of $L^0$. The force directions are indicated by black arrows, and the force-attaching points in the native state and in the transition state are indicated by red dots.

##### XIII. Estimation of the physiologically relevant tension range in integrin-mediated mechanosensing supramolecular linkage

The extension of a mechanosensing supramolecular linkage during cytoskeleton contraction leads to a time varying tension in the linkage, which can be divided into three stages. In the initial stage, low tension stage where all the tension-bearing domains are folded and thus a tighter linkage, the tension increases at a faster rate. In the second stage where the tension-bearing domains in the linkage undergo tension-dependent unfolding transitions, the tension is buffered within a narrow range of several pN [14–16] and hence a low rate of tension increase. In the third stage when all the tension bearing domains unfold, the tension may increase at faster rate till the breakage of the linkage at various tension-bearing protein-protein connections. Overall, the rate of the tension increase (i.e., the loading rate) can be roughly estimated based on the dynamic range of extension in a mechanosensing process. For example, during testing of the ECM stiffness, the cell leading edge protrudes and contracts with a displacement on the order of a few hundreds of nanometers [17]. These oscillations occur with a typical timescale of tens of seconds, and the tensions through individual tension-transmission linkages at focal adhesions are typically between approximately 2 and 10 pN within the tension-buffered stage [18, 19]. Thus, the physiological pulling rate is on the order of tens of nm/s and the force-loading rate on the order of a few pN/s. Figure S6 shows the rupturing tension distribution of the three TGTs under tensions of different loading rates from 1 pN to 10 pN. With the same stretching constraint (loading rate), longer TGTs with higher mechanical stability can sustain for longer time and can potentially enable more effective mechanotransduction.

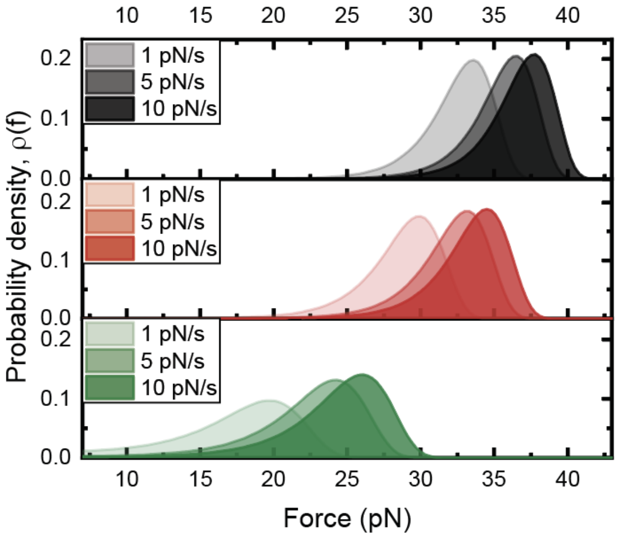


FIG. S6. The rupturing tension probability densities of TGTs when the tension increases at constant loading rates from 1 pN to 10 pN for 15-bp, 13-bp, and 11-bp TGT from top to bottom.


[1] X. Zhao, X. Zeng, C. Lu, and J. Yan, Studying the mechanical responses of proteins using magnetic tweezers, Nanotechnology **28**, 414002 (2017).

[2] H. Chen, H. Fu, X. Zhu, P. Cong, F. Nakamura, and J. Yan, Improved high-force magnetic tweezers for stretching and refolding of proteins and short dna, Biophysical journal **100**, 517 (2011).

[3] S. B. Smith, Y. Cui, and C. Bustamante, Overstretching b-dna: the elastic response of individual double-stranded and single-stranded dna molecules, Science **271**, 795 (1996).

[4] A. Bosco, J. Camunas-Soler, and F. Ritort, Elastic properties and secondary structure formation of single-stranded dna at monovalent and divalent salt conditions, Nucleic acids research **42**, 2064 (2014).

[5] S. Guo, A. K. Efremov, and J. Yan, Understanding the catch-bond kinetics of biomolecules on a one-dimensional energy landscape, Communications Chemistry **2**, 1 (2019).

[6] X. Wang and T. Ha, Defining single molecular forces required to activate integrin and notch signaling, Science **340**, 991 (2013).

[7] J. SantaLucia, H. T. Allawi, and P. A. Seneviratne, Improved nearest-neighbor parameters for predicting dna duplex stability, Biochemistry **35**, 3555 (1996).

[8] J. M. Huguet, M. Ribezzi-Crivellari, C. V. Bizarro, and F. Ritort, Derivation of nearest-neighbor dna parameters in magnesium from single molecule experiments, Nucleic acids research **45**, 12921 (2017).

[9] H. A. Kramers, Brownian motion in a field of force and the diffusion model of chemical reactions, Physica **7**, 284 (1940).

[10] J. SantaLucia, A unified view of polymer, dumbbell, and oligonucleotide dna nearest-neighbor thermodynamics, Proceedings of the National Academy of Sciences **95**, 1460 (1998).

[11] S. Guo, Q. Tang, M. Yao, H. You, S. Le, H. Chen, and J. Yan, Structural–elastic determination of the force-dependent transition rate of biomolecules, Chemical science **9**, 5871 (2018).

[12] J. R. Wenner, M. C. Williams, I. Rouzina, and V. A. Bloomfield, Salt dependence of the elasticity and overstretching transition of single dna molecules, Biophysical journal **82**, 3160 (2002).

[13] X. Zhang, H. Chen, S. Le, I. Rouzina, P. S. Doyle, and J. Yan, Revealing the competition between peeled ssdna, melting bubbles, and s-dna during dna overstretching by single-molecule calorimetry, Proceedings of the National Academy of Sciences **110**, 3865 (2013).

[14] M. Yao, B. T. Goult, B. Klapholz, X. Hu, C. P. Toseland, Y. Guo, P. Cong, M. P. Sheetz, and J. Yan, The mechanical response of talin, Nature communications **7**, 1 (2016).

[15] S. Le, X. Hu, M. Yao, H. Chen, M. Yu, X. Xu, N. Nakazawa, F. M. Margadant, M. P. Sheetz, and J. Yan, Mechanotransmission and mechanosensing of human alpha-actinin 1, Cell reports **21**, 2714 (2017).

[16] M. Yu, Z. Zhao, Z. Chen, S. Le, and J. Yan, Modulating mechanical stability of heterodimerization between engineered orthogonal helical domains, Nature communications **11**, 1 (2020).

[17] G. Giannone, B. J. Dubin-Thaler, H.-G. Döbereiner, N. Kieffer, A. R. Bresnick, and M. P. Sheetz, Periodic lamellipodial contractions correlate with rearward actin waves, Cell **116**, 431 (2004).

[18] C. Grashoff, B. D. Hoffman, M. D. Brenner, R. Zhou, M. Parsons, M. T. Yang, M. A. McLean, S. G. Sligar, C. S. Chen, T. Ha, *et al.*, Measuring mechanical tension across vinculin reveals regulation of focal adhesion dynamics, Nature **466**, 263 (2010).

[19] K. Austen, P. Ringer, A. Mehlich, A. Chrostek-Grashoff, C. Kluger, C. Klingner, B. Sabass, R. Zent, M. Rief, and C. Grashoff, Extracellular rigidity sensing by talin isoform-specific mechanical linkages, Nature cell biology **17**, 1597 (2015).
